# FT-Kernel: An innovative kernel for decoding cellular secrets related time

**DOI:** 10.1101/2025.03.01.640966

**Authors:** Cencan Xing, Yuqing Ma, Yanting Wang, Yixuan Wang, Peiwu Qin, Hongwu Du, Zehua Zeng

## Abstract

The cell fate participants characterization based on single-cell RNA sequencing (scRNA-seq) data greatly facilitates the mechanism understandings of cellular differentiation. However, inferring these fate factors dynamics along the pseudotime is challenging. Based on cell-state density and pseudotime regression weights, we present an algorithm TimeFactorKernel (FT-Kernel), to predict the key cell fate factors, not only the minimum lineage transition genes, but also the related genesets/pathways, and cellular interaction dynamics along the pseudotime. By extrapolating the pseudotime-related key genes from spectral data as a pseudotime-kernel, FT-Kernel outperformed previous methods. Beyond time-related genes, FT-Kernel offered a comprehensive analysis of the inferred cellular interactions dynamics and lineage pathways dynamics along the pseudotime, which were limited in other methods. Additionally, it facilitated time-related fate factors prediction across various data modalities. Our work developed an important pseudotime-kernel in predicting the fate factors, and provided insights into the cellular hierarchies during development.

**Significance:** Understanding cellular changes over time is key to deciphering life’s processes from development to disease. Current single-cell methods miss crucial information by overlooking cell-state density dynamics and thus neglecting rare, low-density transitional states vital for cell fate decisions. TimeFactorKernel, a novel computational tool, overcomes this limitation. By prioritizing genes active in these low-density transitional states and employing machine learning for precise gene identification, TimeFactorKernel provides a powerful new approach to decode cell fate. Validated and widely adopted, TimeFactorKernel also uniquely models cell-cell interactions, opening new avenues for understanding complex biological systems in health and disease.

**Keypoints:**

- FT-Kernel is a novel kernel algorithm designed to predict time-related fate factors dynamics from the cell state density and pseudotime regression weights.
- FT-Kernel can be applied to identify lineage transition key genes, cellular interactions, and the multimodal fate kernel inference of genesets/pathways.
- Compared to other temporal-related kernels, FT-Kernel minimizes the skewness of gene distribution and more accurately captures the state of fate transitions.
- FT-Kernel is an extensible multimodal framework facilitating to understand the mechanisms of cell differentiation and is available on GitHub.

## Introduction

Differentiation is the process by which cells develop into different types, and it’s essential for the growth and function of all multicellular organisms^1^. Understanding the regulators that control this process is crucial for grasping how development works and what goes wrong in diseases^2,3^. With the development of single-cell RNA sequencing (scRNA-seq), scientists can now use methods like cell trajectory analysis and cell-state density estimation to study how cells change over time^4,5^. These methods use data from individual cells to classify them into different developmental paths, calculate a timeline called pseudotime^6^, and identify where cells are along their pathway. By analyzing pseudotime, we can track how gene expression changes during development and understand the dynamics of these biological processes ^7–9^.

Traditional methods often focus only on the relationship between pseudotime and gene expression when identifying genes that define cell lineages ^10–14^. However, the density information of cell states was usually ignored, which means the complex dynamics within the cell population was thereby missed out. Cell-state density is defined as the local probability density of cells along pseudotime trajectories, computed via Mellon’s Poisson point process^7^. Transitional states (e.g., branching points) correspond to low-density regions (density < 0.5×max lineage density; see Methods) ^15,16^. When a cell type changes, it moves from high-density to low-density and back to high-density areas ^17,18^. This pattern shows how cell states change dynamically in populations. Genes that change significantly between low and high-density states are often linked to cell transformations, but these changes don’t follow a simple timeline ^7^. Additionally, gene activity doesn’t always follow a straight path and gene expressions are often linked, these methods mainly find genes active at the initiate and terminal states with skewness of pseudotime. As a result, they might miss genes that play roles in the middle state of these processes.

Unlike existing tools (e.g., Monocle, Slingshot) that rely on Pearson correlation, FT-Kernel uniquely integrates cell-state density weighting with ridge regression to prioritize transitional-state genes. This dual integration represents the first method to bridge density dynamics with pseudotime analysis. Rather than employing traditional correlation kernels, we apply a regression approach to identify key differentiate factors with time-dynamics for specific lineages through shifts in cell-state density. Given the multicollinearity among factors, we employ ridge regression as the underlying model. Additionally, our inputs are compatible with all commercially available models of proposed temporal sequencing algorithms, fully leveraging the potential of various models to ensure compatibility with scRNA-seq generated by different techniques.

In this study, we commence by demonstrating FT-Kernel’s efficacy in identifying cell-state density lineage transition genes involved in human hematopoiesis and mouse dentate gyrus neuron developmental processes. Subsequently, by applying FT-Kernel to the decidua-placenta interface dataset, it can identify critical cellular interactions during lineage transitions in trophoblast development. Additionally, FT- Kernel could identify cell-state density lineage transition pathways in hematopoiesis. Our results indicate that FT-Kernel provides a novel perspective on the study of genealogical developmental determinant genes/gene sets. FT-Kernel is available as an open-source package at https://github.com/Starlitnightly/FT-Kernel.

## Result

### The FT-Kernel modeling approach

To precisely identify the lineage transition genes related to cell-state density from high-dimensional single-cell phenotypic landscapes, we set to decompose these landscapes into different lineages driven by a series of factors based on the pseudotime and cell-state density (Fig. 1a). Here are two main challenges need to be addressed: 1) First, calculating the gene variability from the cells in low-density regions. Cells in low-density areas often represent rare or transitional states that are biologically important. Traditional methods tend to overlook the changed genes of cells in low-density regions. 2) The second challenge is reducing noise. While linear regression can calculate weights for all genes, analyzing all genes can introduce noise. FT-Kernel addresses this by using a technique called ridge regression to identify important genes (Fig. 1b).

**Fig. 1.**
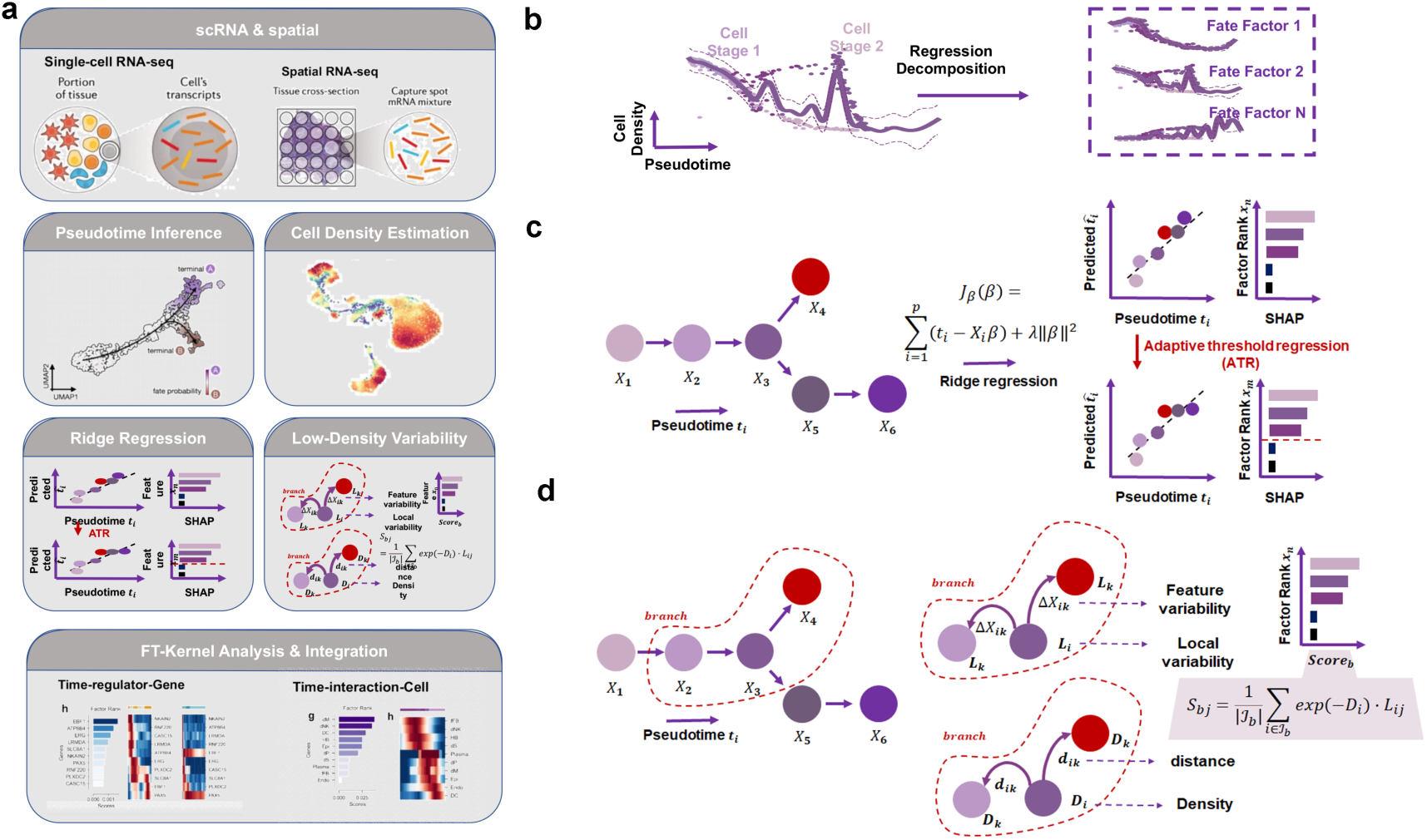
Workflow of the FT-Kernel algorithm. (a) Data Input and Preprocessing. FT- Kernel takes scRNA-seq data as input, and performs pseudotime inference and cell- state density estimation using methods like Palantir and Mellon, respectively. This preprocessing step prepares the data for downstream FT-Kernel analysis. The overall workflow enables the identification of key lineage transition genes, cellular interactions, and fate-determining pathways involved in cell differentiation and development. (b) Decomposition of the density of cell states along the pseudotime. We design to use different sets of time-related genes/fate factors for cell-state density reconstruction. For the density of states of a cell, we are obtaining the neighborhood distances of cells from their gene expression patterns, and then using a Gaussian process to connect the densities between highly similar cell states in order to compute the density of states of a cell characterizing a single-cell phenotypic landscape. (c) The minimum sets of time-related genes prediction. We used 𝑋_!_ to denote the gene expression pattern of each cell and used it as an input to the ridge regression, and 𝑡_!_ to denote the proposed time-series value of each cell, and we subsequently minimised the cost function for fitting the ridge regression model. After obtaining the weights for each gene, the genes are further filtered by adaptive threshold regression (ATR) model with interpreted by the unified framework Shapley additive explanations (SHAP). We calculated the SHAP value for each gene, progressively narrowing down the SHAP threshold and thus obtaining the minimum set of genes with the same regression coefficient of determination. (d) The genesets predication for low-density of cell states. The local gene variability 𝐿_!_ for each cell and for a given special lineage branch 𝑏 is calculated. Then we calculated the low-density variability score for each highly SHAP gene 𝑗, and weighted summed all cells within the lineage to obtain the overall low-density variability score for gene 𝑗 within the lineage.

In this study, the calculations for time-related genes (Fig. 1b) and low-density regions (Fig. 1c) are done independently. For identifying time-related genes, we recommend using the pseudotime of all cells within a lineage to build the ridge regression model (Fig. 1b). To find a minimal set of genes necessary for accurate analysis, FT-Kernel uses a method called adaptive threshold regression (ATR), to acquire the minimum set of genes by progressively narrowing down the shapley additive explanations (SHAP) value, providing a clear view of the essential fate factors for cell lineages development (Fig. 1b).

For calculating low-density regions of cells (Fig. 1c), we introduced the Mellon method ^7^. We first calculate the distance from each cell to its nearest neighbor and use a Poisson point process to relate this distance to local density. Higher density areas correspond to closer neighbor distributions, while narrower density areas have wider distributions. With this density pattern, Branch selection follows a biologically informed heuristic approach: (1) Perform Leiden clustering on all cells. (2) Manually select clusters spanning transitional states (e.g., clusters containing both progenitor and differentiated cells). (3) Validate branches by ensuring they include branching origins (cells shared by multiple lineages). This process balances biological interpretability with computational simplicity. If we only use pseudotime from the pre-developmental branch, we might find more “local time-related genes,” but these could also show “local temporal” characteristics elsewhere in the lineage. FT-Kernel then calculates the variability of each gene in each cell within these branches and weighted sums them with time-related genes all to get the overall variability within the branch.

Interestingly, FT-Kernel can measure not only genes within a lineage along the pseudotime, but also the gene sets within a lineage, cellular interactions, and lineage determined pathways, which analyses are not compatible with velocity-processed datasets. This means that FT-Kernel can reveal these time-relate factors at the micro level, from lineage transition genes, lineage pathways, to the cellular interactions, in addition to the temporal nature of cellular regulation and function at the overall level.

### Kernel benchmarking of genetic algorithms with different pseudotime correlations

To benchmark FT-Kernel against previous methods for identifying pseudotime-related genes, we first examined three test dataset commonly used in pesudotime analysis, the mouse pancreas developmental data (Fig. 1), mouse dentate gyrus neuron (DG) developmental data (Fig. S1a-c), and human hematopoietic stem cells (HSC) developmental data (Fig. S1d-f). To obtain the pseudotime, for velocity-based data, we used dynamic model scVelo to compute latent time as pseudotime (Fig. 1b and Fig. S1b); and for data not based on velocity, we used palantir to compute pesudotime (Fig. S1e). Then we used the pancreatic differentiation profiles in mice to compare three models, the traditional Pearson correlation coefficient (Pearsonr), Gene Likelihoods (only available for velocity-based data), and FT-Kernel (Fig. 2c-h). We found that the pseudotime-related genes identified by Pearsonr exhibited a clear distribution skewness for pseudotime distribution of only maxima or minima without median, while those identified by Gene Likelihood and FT-Kernel showed a continuous distribution along the pseudotime and no pseudotime distribution skewness, which means the sorting growth of pseudotime is linear, not jumpy (Fig. 2c, Fig. S1c, and Fig. S1f). Following benchmarking standards in pseudotime analysis^19^, we selected R² for regression accuracy and Kendall’s τ for temporal ordering consistency. Furthermore, we used the regression coefficient of determination to characterize the temporal distribution of pseudotime-related genes found by these different models (Fig. 2d). The Pearsonr model had the lowest regressivity with a coefficient of 0.88, while FT-Kernel reached a similar level as Gene Likelihoods. Note that FT-Kernel and Pearsonr can model any arbitrary pseudotimes datasets, whereas Gene Likelihoods is limited to scVelo’s dynamic model. To comprehensively evaluate the performance of FT-Kernel in identifying pseudotime-related genes, we selected a panel of metrics encompassing different aspects of performance. Number of Genes was chosen to assess the model’s ability to identify a minimal set of genes, reflecting a balance between capturing temporal dynamics and minimizing noise and experimental cost. R-squared (R²) and Mean Absolute Error (MAE) were used to evaluate the regression accuracy of pseudotime prediction using the identified gene sets, ensuring that the selected genes retain pseudotime information comparable to traditional methods like Pearson correlation. Kendall’s tau was included to measure the temporal ordering consistency of the identified genes, reflecting the model’s ability to capture the temporal progression of gene expression. Furthermore, we introduced Linregress slope as a novel metric to assess the linearity of gene expression changes along pseudotime, addressing the limitations of Pearson correlation in capturing genes active in intermediate states.

**Fig. 2.**
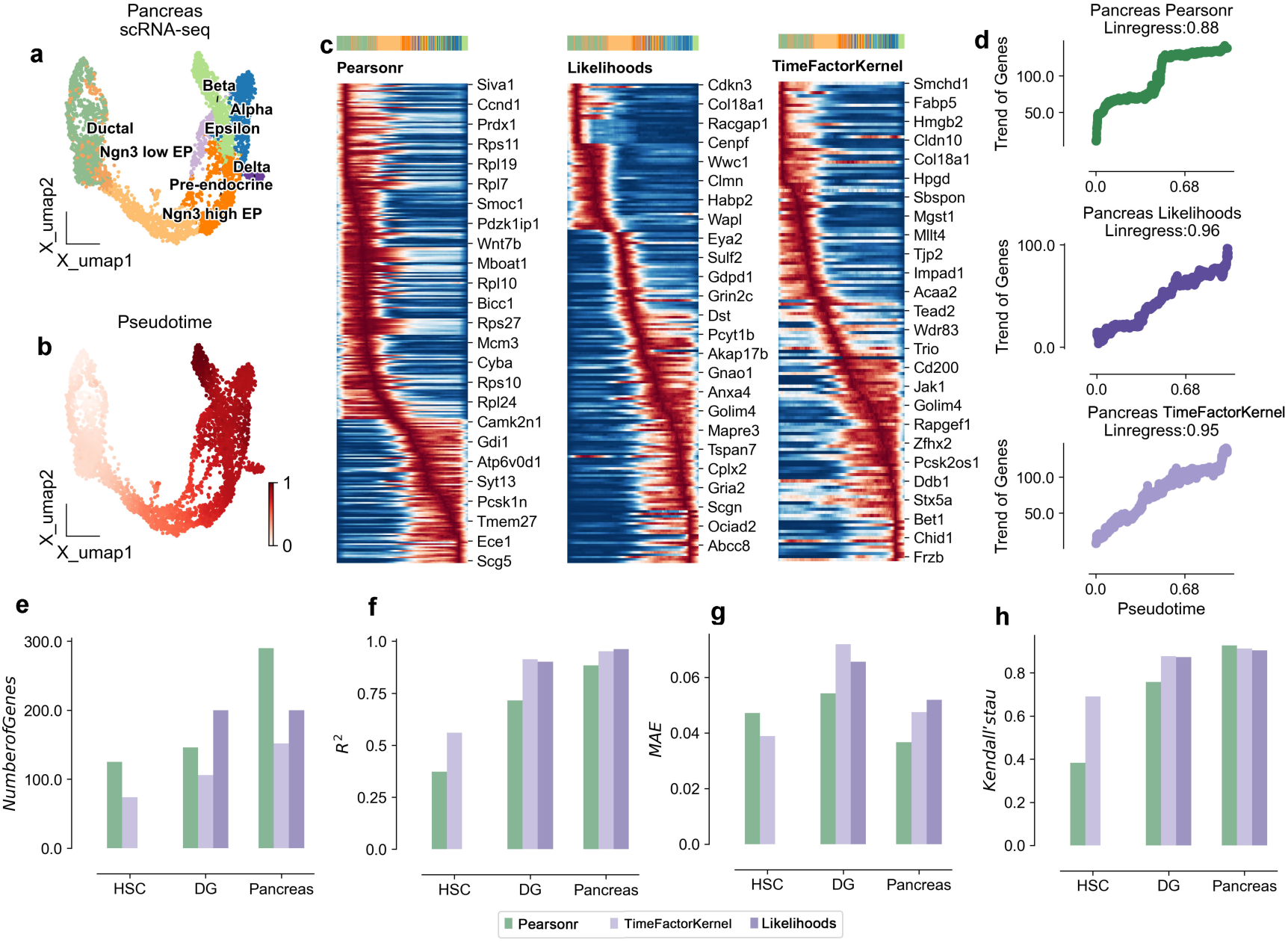
Kernel benchmarking test of FT-Kernel. (a) UMAP of mouse pancreas development scRNA-seq data^19^, colored by cell types. (b) Same as a, colored by psdutotime calculated by scVelo dynamic model. (c) Heatmap of pseudotime related genes, from left to right the calculations are analyzed by Pearsonr, Gene Likelihoods, and FT-Kernel respectively. (d) Distribution of expression regions of pseudotime related genes calculated by three different algorithms. The coefficient values were shown for each algorithm. (e) Comparison of pseudotime-related numbers of three different algorithms on human hematopoietic stem cells (HSC), mouse dentate gyrus neurons (DG) and mouse pancreas development, respectively. (f) Comparison of regression correlation coefficient. (g) Comparison of Regression Mean square error (MAE). (h) Comparison of Kendall’s tau correlation of pseudotime-related genes in the trajectory.

Next, we further used the other three metrics to assess how FT-Kernel differs from other models on different data. We first examined Number of Genes, a common metric for evaluating the discovery of pseudotime-related genes; a larger number of genes implies the introduction of more timing-independent noise. For the mentioned HSC, DG, and Pancreas datasets, FT-Kernel identified the fewest Number of Genes (Fig. 2e, since HSC datasets were not computed by scVelo, Gene Likelihoods could not apply on HSC datasets), but these genes retained the characteristics of temporal regression (Fig. 2d). We then evaluated the coefficient of determination (R^2) and Mean Absolute Error (MAE), which are regression metric for assessing pseudotime-related genes. We performed linear regression modeling using the pseudotime-related genes identified by the models and evaluated the ratio of the variance of the explanatory variables of the proposed time series predicted by the regression model to the true time series. For the HSC, DG, and Pancreas datasets, the R^2 of FT-Kernel was similar to that of Gene Likelihoods and significantly higher than that of Pearsonr (Fig. 2f). Additionally, the MAE did not exceed an error of 0.01 across the three models (Fig. 2g). We also assessed Kendall’s tau, a metric for the temporal ranking of pseudotime-related genes. We calculated the mean position of each cell with greater than 80% gene expression and measured that position (see Methods for details), which in turn evaluates the consistency of the cell’s high-temporal mean position with the actual. FT-Kernel and Gene Likelihoods exhibited the highest concordance of cellular temporal ordering of genes (Fig. 2h). Considering the limitation of Gene Likelihoods in analyzing the non-velocity-based datasets, FT-Kernel can apply to arbitrary pseudotimes datasets, and exhibit a well performance as Gene Likelihoods.

### FT-Kernel resolves cellular development density lineage transition genes

Considering FT-Kernel has found the fewest pseudotime-related genes, we set to examine whether these genes are closely linked to the fate of genealogical development. We first explored the potential of FT-Kernel using human hematopoietic stem cell (HSC) differentiation lineage data. This dataset was divided into three different differentiation directions, B-cell fate profiles, erythroid fate profiles, and monocyte fate profiles (Fig. 3a). We calculated pseudotime using Palantir and cell- state density using mellon (Fig. 3b-3c). FT-Kernel captures pseudotime-related genes across the entire spectrum of HSC fates (Fig. 3d), especially in the median pseudotime region with lowest cell-state density (Fig. 3e). We also used Leiden unsupervised clustering method, identifying 23 cell clusters within the entire HSC fate lineage (Fig. 3f).

**Fig. 3.**
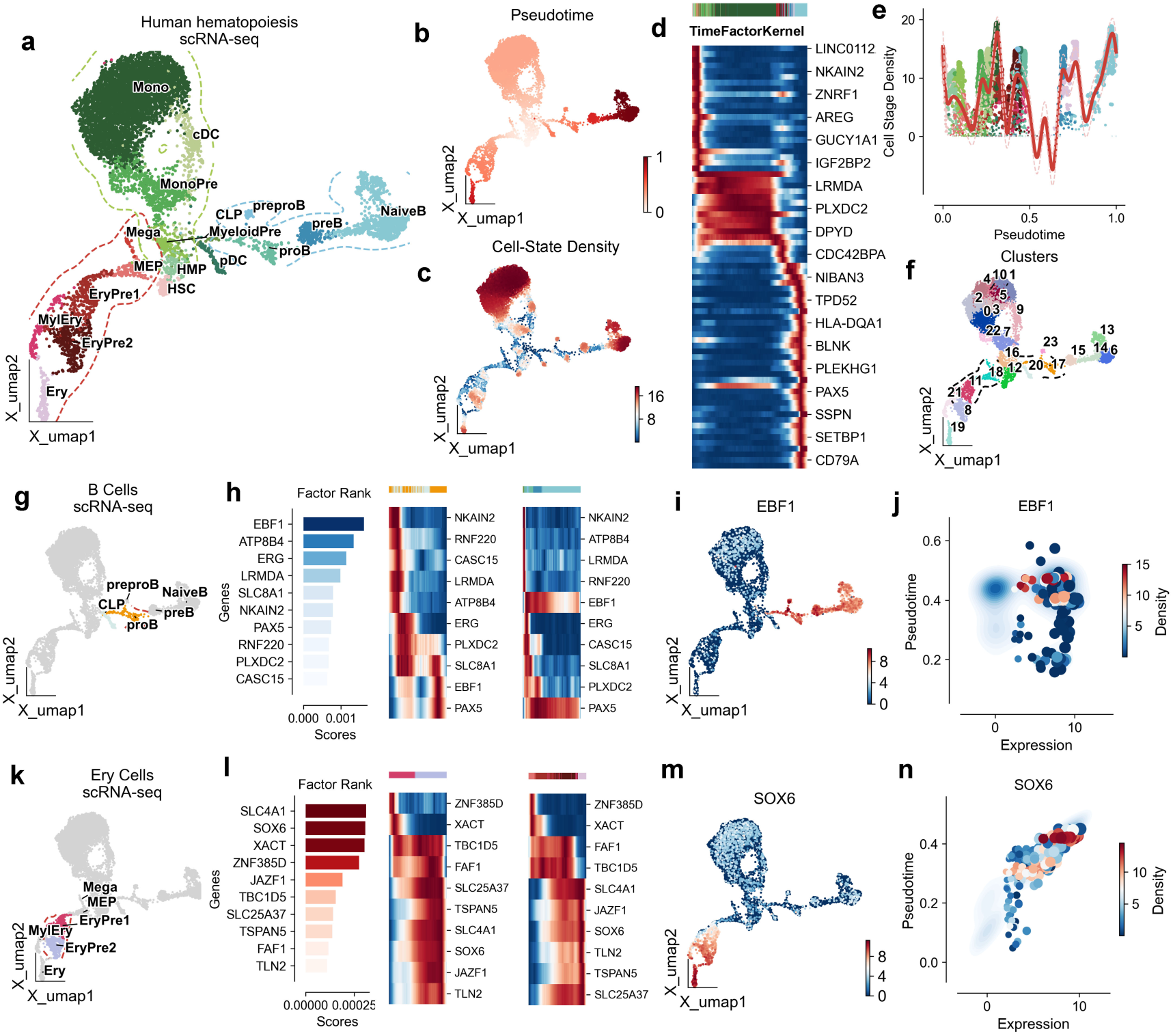
FT-Kernel reveal the lineage determination genes in human hematopoiesis. (a) UMAP of T-cell-depleted bone marrow scRNA-seq data^26^, colored by cell types. (b) Same as a, colored by pseudotime calculated by Palantir. (c) Same as a, colored by cell-state density calculated by Mellon. (d) Heatmap of pseudotime related genes calculated by FT-Kernel. (e) Comparison of pseudotime and log density for hematopoietic lineages. Cells highlighted in all cell types. (f) Same as a, colored by unsupervised clustering by Leiden. (g) Cells from B-cell lineages, tracing differentiation from early stages and colored by Leiden cluster. (h) The top 10 cell-state density lineage transition genes identified by FT-Kernel, from left to right are rank plots, heatmaps of genes changing over pseudotime in the selected Leiden Clusters, and heatmaps of genes changing over pseudotime in the B-Cells lineage. (i) UMAPs colored by *EBF1* expression. (j) Cell-state density distribution plot of *EBF1*, horizontal coordinates represent gene expression, vertical coordinates represent proposed pseudotime values, and colored by cellular density. (k) Cells from erythroid lineages, colored by Leiden cluster. (l) The top 10 cell-state density lineage transition genes of erythroid lineages. (m) UMAPs colored by *SOX6* expression. (n) Cell-state density distribution plot of *SOX6*.

For the B-cell differentiation lineage, we selected the clusters ‘17, 20, 23’ with the lowest average timings for the prediction of fate factors in early lineage development (Fig. 3g). FT-Kernel identified 10 regulatory genes within the early lineage based on cell-state density transition, among which *EBF1*, a major regulator of B-cell development ^20^, was ranked highest by FT-Kernel and showed a clear shift in the early lineage (left panel of Fig. 3h). Although a two-wave expression of *EBF1* was shown along the pseudotime of the Leiden unsupervised clusters (middle panel of Fig. 3h), in the overall B cell lineage, *EBF1*’s expression peak in low-density regions aligns with its known role in destabilizing B-cell progenitors^20^ (right panel of Fig. 3h, Fig. 3i). Notably, the high-expression levels of *EBF1* with low-pseudotime regions corresponded to the cells with lower cell-state density (Fig. 3j), indicating that a transitional role of *EBF1* expressed in the lower cell-state density for the B-cell lineage differentiation. *ATP8B4* is a myeloid-specific flippase^21^, and past reports have suggested that the flippase *ATP11C* during early B-cell development^22^. The combination of a high TM-score (0.85) and a reasonably low RMSD (2.85 Å) from the US-align analysis strongly indicates that ATP8B4 and ATP11C are structurally very similar. This structural similarity reinforces the prediction that ATP8B4 functions as a flippase, similar to ATP11C, and supports its role in myeloid cell development. In addition, *EGR* ^23^ and *PAX5* ^24^ with early-stage shift expression were also regulators of B-cell development, further supporting the precise prediction for time-related genes by FT-Kernel.

For the erythroid differentiation lineage, we selected the clusters ‘8, 11, 18, 21’ with the lowest average timings for the prediction of fate factors in early lineage development (Fig. 3k). FT-Kernel identified 10 regulatory genes within the early lineage (left panel of Fig. 3l). The top 4 genes included the erythrocyte membrane protein *SLC4A1*, a major regulator of erythroid development *SOX6* ^25^, lncRNA *XACT*, and ZNF385D, in which the latter two genes exhibited a cute shift in the early lineage (Fig. 3l), suggesting a potential role during erythroid development. *SOX6* showed an elegant activation in the early lineage and sustained high expression in the mid-late lineage throughout the erythroid development (Fig. 3l, 3m). Additionally, *SOX6* expression regions at low pseudotime had lower cell-state density (Fig. 3n), further supporting a more precisely time-related factor prediction of FT-Kernel based on cell- state density.

### Identification of lineage determined cellular interactions dynamics during trophoblast development

In addition to identifying time-related regulatory genes for lineage development, FT-Kernel can also probe regulatory cells and targets of action for lineage development. We selected the maternal-fetal interface as our study object (Fig. 4a), where trophoblast cells invade the uterus. This process is not only regulated by the trophoblast cells themselves but also significantly influenced by the microenvironment of the maternal-fetal interface. We calculated the meta-cells of the maternal-fetal interface using SEACells to reduce data sparsity and improve gene robustness, allowing us to investigate the differentiation of the trophoblast cells. The trophoblast cells were composed with three main epithelial cells: the villi cytotrophoblast (VCT), syncytiotrophoblast (SCT) and extravillous trophoblast (EVT) (Fig. 4b), which then were calculated the cell-state density of the localized trophoblast cells using Mellon (Fig. 4c). We found a trough region mainly composed of SCT cells during the differentiation from VCT to EVT (Fig. 4d). To understand cellular interactions at the maternal-fetal interface, we used CellPhoneDB to calculate interactions involving trophoblast cells as receptor cells, and found that the endothelial cells, epithelial cells, and decidual stromal cells (dS) cells exhibited the most ligand-receptor pairs with trophoblast cells (Fig. 4e-4f).

**Fig. 4.**
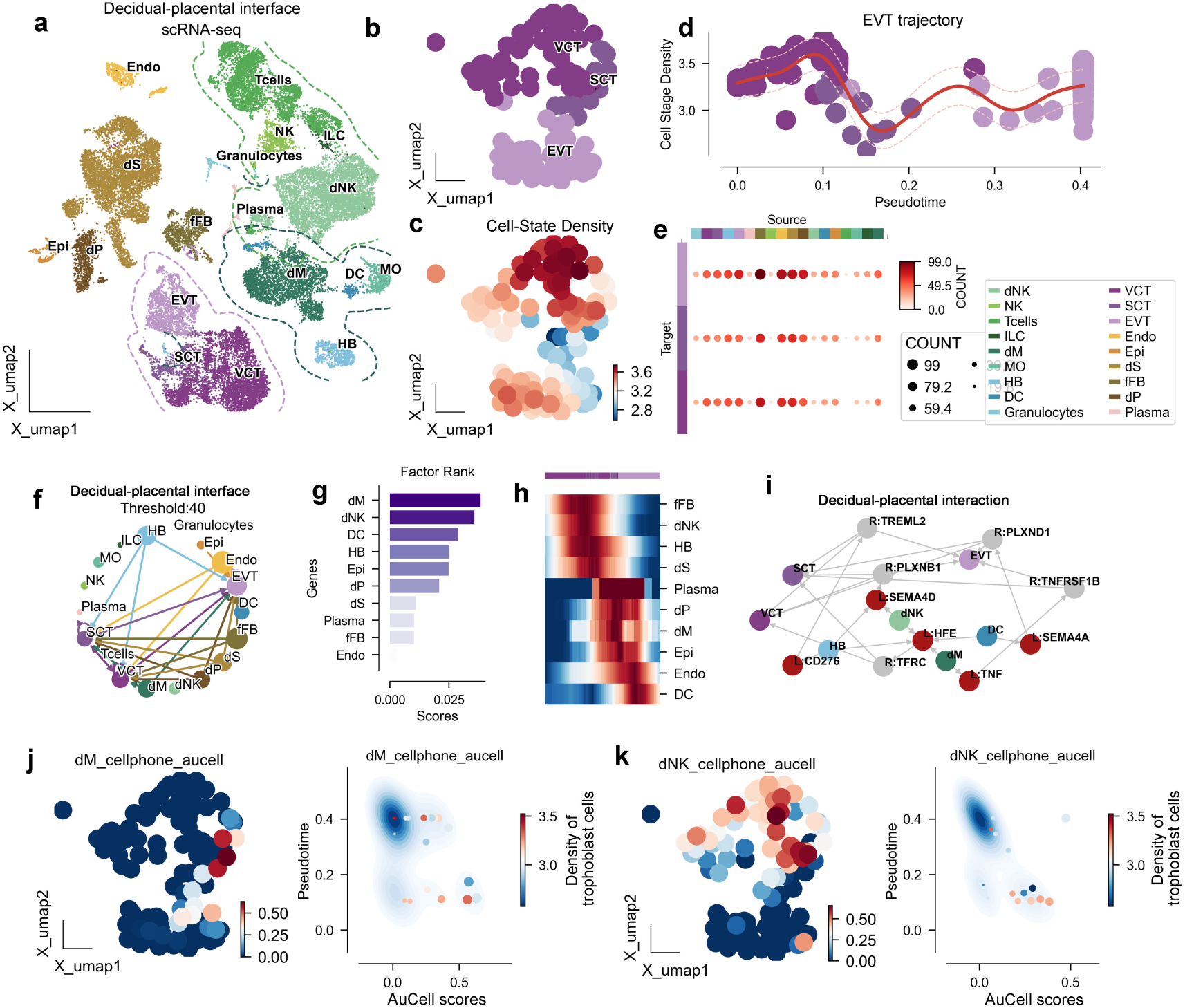
Lineage determined cellular interaction in trophoblast development. (a) UMAP of decidual-placental interface scRNA-seq data ^27^, colored by cell types. (b) UMAP of extravillous trophoblast (EVT) development meta-cells, colored by cell types. (c) Same as b, colored by cell-state density. (d) Comparison of pseudotime and log density for EVT lineage development, colored by cell types. (e) Heatmap showing the cellular interaction between EVT lineage cells and other decidual-placental interface cells. (f) Network showing the cellular interaction targeting EVT lineage cells. (g) Top 10 density-of-state spectral shifts in cellular interactions targeting EVT lineage calculated by FT-Kernel. (h) Heatmap showing the cellular interactions along the pseudotime of EVT lineage. (i) Source-target network of EVT lineage and other decidual-placental interface cells. Except VCT, SCT, and EVT cells representing as ligands, the cells labeled in red also represent ligands; while the cells labeled in grey represent receptors. (j) EVT cells with AUCell scores targeted by dM cells and cell-state density. (Left) Same as b, colored by the AUCell scores of receptors on EVT lineage cells targeted by dM cells. (Right) Cell-state density distribution plot of EVT lineage cells with dM AUCells scroes, horizontal coordinates represent AUCell score, vertical coordinates represent proposed pseudotime values, and colored by cellular density. (k) EVT cells with AUCell scores targeted by dNK cells and cell-state density. (Left) Same as b, colored by the AUCell scores of receptors on EVT lineage cells targeted by dNK cells. (Right) Cell-state density distribution plot of EVT lineage cells with dNK AUCells scroes. dNK, decidual nature killer cells; NK, nature killer cells; ILC, innate lymphocyte cells; dM, decidual macrophages; MO, monocytes; HB, Hofbauer cells; DC, dendritic cells; VCT, villous cytotrophoblast; SCT, syncytiotrophoblast; EVT, extravillous trophoblast; Endo, endothelial; Epi, epithelial; dS, decidual stromal cells; fFB, fibroblasts; dP, decidual perivascular cells.

Next, we selected the uniquely paired receptor on trophoblast cells for each cell class as regulators targeting trophoblast cells. We used the AUCell scores of these receptors on trophoblast cells for modeling and as inputs to FT-Kernel. The highest fate-regulated cells identified by FT-Kernel were decidual macrophages (dM) and decidual nature killer cells (dNK) (Fig. 4g). The dM and dNK regulatory time points were located at mid-late (during VCT-SCT-EVT transition) and early (within VCT) stages of trophoblast differentiation, respectively (Fig. 4h). For dM, they mainly target the receptors *TFRC* and *TNFRSF1B* on trophoblast cells via *HFE* and *TNF* (Fig. 4i). Trophoblast cells with high AUCell scores interacting with dM cells are predominantly located in the SCT region (left panel of Fig. 4j), and the density of trophoblast cells is located in an intermediate state in the region with high AUCell scores and low temporal order (Fig. 4j). For dNK, they target receptors *TFRC* and *PLXNB1* on trophoblast cells via *HFE* and *SEMA4D* (Fig. 4i). Areas with high AUCell scores for dNK are mainly located in the VCT and early SCT, with trophoblast cell-state density also in an intermediate state in high AUCell scores and low temporal order regions (Fig. 4k). Interestingly, the developmental effects of dNK and dM on trophoblast cells have been previously reported ^27^. Our algorithm revealed the critical regulatory dynamics of dNK and dM on trophoblast cell development from a data perspective.

### Identification of pathways dynamics during lineage determined process in hematopoiesis

Finally, we demonstrate the ability of FT-Kernel in recognizing pseudotime-related functions, which in past studies have generally been identified using Pearsonr, followed by pathway enrichment of the pseudotime-related genes. This method relies on pseudotime-related genes and assumes that these genes do not overlap across different temporal modules. However, developmental processes may have a continuous role for a particular gene, such as the B-cell lineage regulator *PAX5* identified before (Fig. 3h), suggesting that we should quantify the genesets/pathways as a whole.

We still selected human hematopoietic stem cell (HSC) lineage and used Biological Process from Gene Ontology as a pathway database. We used AUCell to score 6036 genesets for all cells from HSC. We then modeled ridge regression on pseudotime using these 6036 genesets, achieving a regression coefficient of 0.94 (Fig. 5a). By maintaining the regression cell fluctuations within ±0.02, we obtained the minimum genesets using FT-Kernel’s ATR, comprising a total of 564 genesets (Fig. 5b-5c). We analyzed the local variability of these time-related genesets using Mellon (Fig. 5d).

**Fig. 5.**
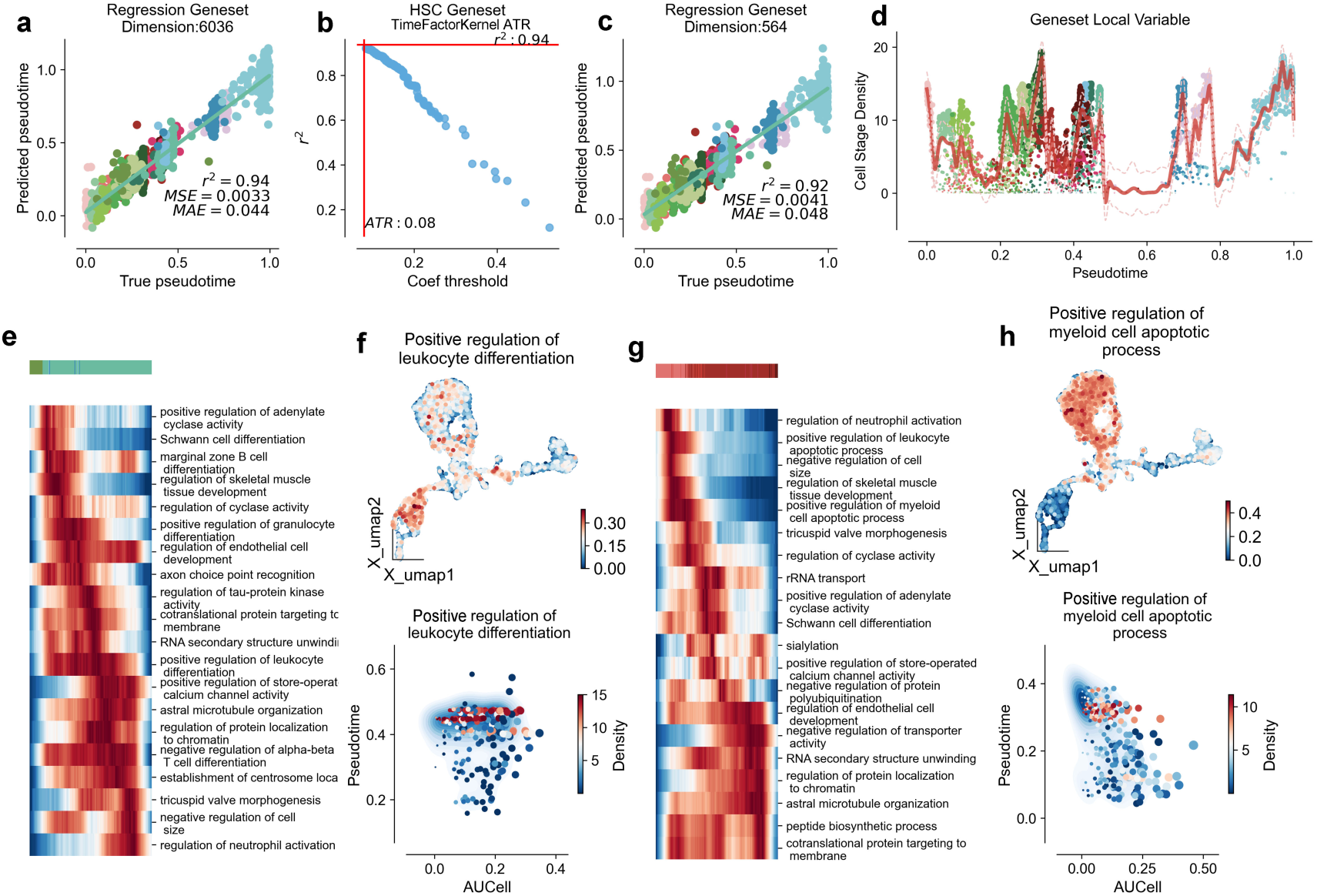
Lineage determined pathways in hematopoiesis development. (a) Pseudotime regression using all 6036 pathway of Gene Ontology biological. (b) Calculation of adaptive threshold regression (ATR) by FT-Kernel reducing the range of pseudotime- related pathways. (c) Pseudotime regression using 564 pathway. (d) Comparison of pseudotime and log density for hematopiuetic lineages. Cells were highlighted in all cell types. (e) Heatmap showing the cell-state density lineage transition pathways analyzed by FT-Kernel in B cells lineage. (f) (Upper) UMAPs of T-cell-depleted bone marrow scRNA-seq data, colored by AUCell scores of positive regulation of leukocyte differentiation. (Bottom) Cell-state density distribution plot of positive regulation of leukocyte differentiation with AUCell scores, horizontal coordinates represent AUCell scores, vertical coordinates represent proposed pseudotime values, and colored by cellular density. (g) Heatmap showing the density lineage transition pathway by FT-Kernel in Erythroid lineage. (h) (Upper) Same as f, colored by AUCell score of positive regulation of myeloid cell apoptotic process. (Bottom) Cell-state density distribution plot of positive regulation of myeloid cell apoptotic process AUCells.

Then we first assessed the temporal correlation genesets in B-cell differentiation profiles, and obtained the top 20 time-related pathways with the highest local variability based on cell-state density from the above 564 pathways (Fig. 5e). Among which we captured the positive regulation of leukocyte differentiation (GO: 1902107), with a continuous activation except the initiate and terminal stage during B-cell lineage development (Fig. 5e). The high AUCell scores distribution of the positive regulation of leukocyte differentiation mainly located in the B-cell and erythroid lineages (upper panel of Fig. 5f), and similarly, the low-pseudotime regions corresponded to the cells with lower cell-state density (bottom panel of Fig. 5f), suggesting an important role of this pathway at the early stage of B-cell lineage.

Next, we evaluated the temporal correlation genesets in the erythroid differentiation lineages and selected the top 20 pathways with the highest local variability (Fig. 5g), among which we captured positive regulation of myeloid cell apoptotic process (GO:0033034) showing a clear shift at the early stage of erythroid differentiation lineages (Fig. 5g, and upper panel of 5h), where HSC differentiate into erythrocytes through a process known as erythroid differentiation. During the differentiation of hematopoietic stem cells into erythrocytes, they initially pass through a cellular stage known as the “myeloid cell lineage”. The apoptosis of myeloid lineage cells is part of the regulation of erythropoiesis and maintenance of homeostasis, the low-pseudotime regions of high AUCell scores corresponded to the cells with lower cell-state density (bottom panel of Fig. 5h), indicating an adjustment of erythroid maintenance of homeostasis at the early stage.

### FT-Kernel Accurately Models Pseudotime and Elucidates Key Fate Determinants in Human Limb Bud Development

To rigorously assess the versatility and robustness of FT-Kernel, we extended its application to a recently published, spatially resolved single-cell transcriptomic atlas of human embryonic limb bud development. This intricate dataset meticulously maps cellular states in both spatial and temporal dimensions during limb morphogenesis^28^, providing an exceptional platform to evaluate FT-Kernel’s performance in a spatially contextualized developmental process. We performed pseudotime inference and cell-state density estimation on the limb bud data, subsequently employing FT-Kernel to dissect the critical fate determinants orchestrating limb bud development.

Consistent with the original spatial annotation^19^ UMAP visualization of limb bud cells, colored by regional identity, distinctly demarcated spatial domains corresponding to ‘Digit’, ‘DistMes’, ‘HandPlate’, ‘IDS’, and ‘Other’ compartments (Fig. 6a). Pseudotime analysis effectively recapitulated the developmental trajectory within the limb bud, as visualized within the UMAP embedding (Fig. 6b). Cell-state density estimation further resolved transitional cellular states integral to the limb bud lineage progression (Fig. 6c). Application of FT-Kernel to this spatially resolved dataset revealed a high degree of concordance between FT-Kernel predicted pseudotime and the inferred developmental timeline, evidenced by a low Root Mean Square Error (RMSE) of 0.088 (Fig. 6g). This quantitative metric underscores FT-Kernel’s accuracy in capturing the intrinsic temporal dynamics of limb bud morphogenesis.

**Fig. 6.**
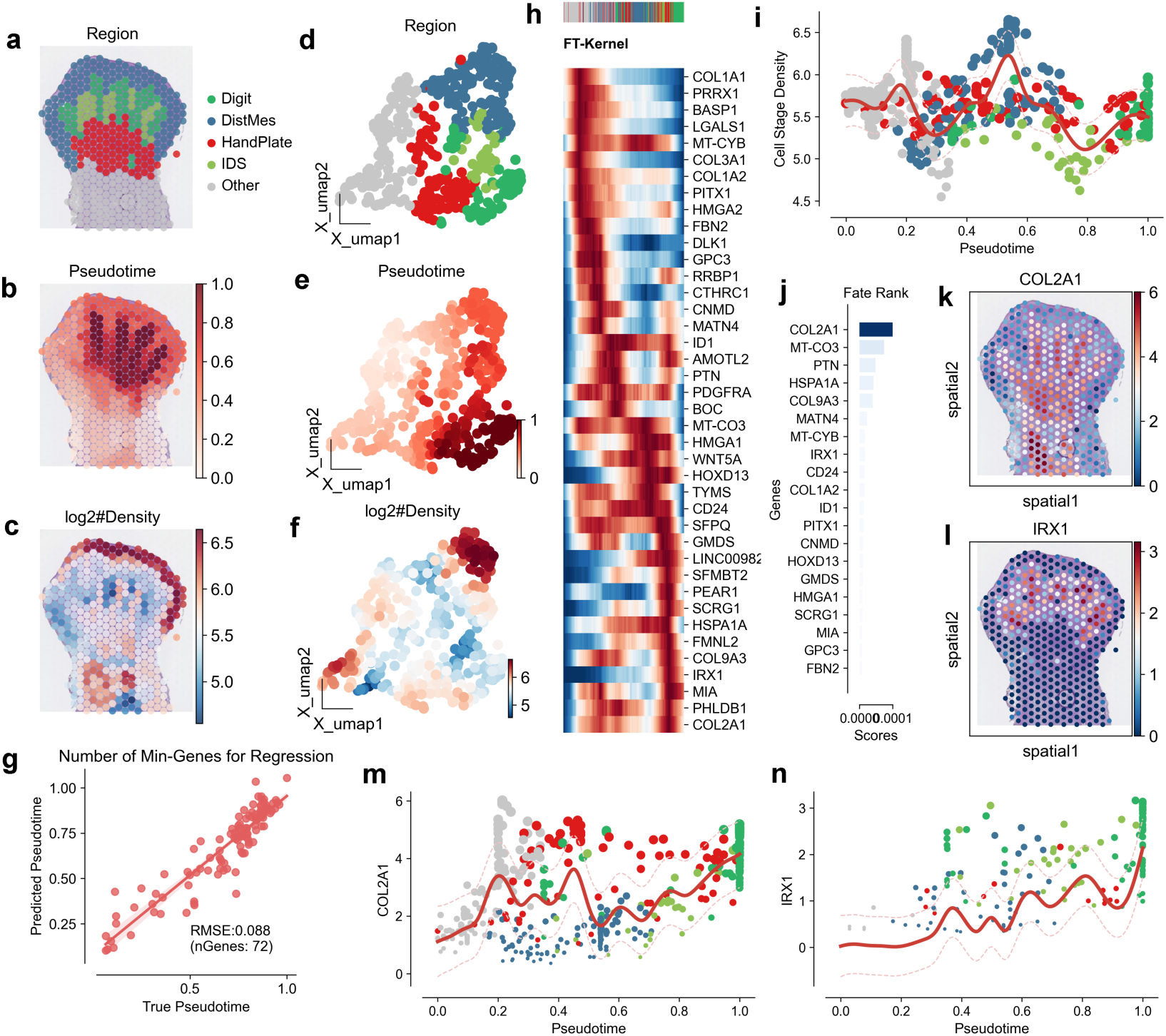
FT-Kernel Analysis of Human Embryonic Limb Bud Development Uncovers Key Fate Determinants. (a) UMAP visualization of human embryonic limb bud scRNA-seq data, colored by annotated anatomical regions of the developing limb. (b) UMAP visualization of limb bud cells, colored by pseudotime inferred by FT-Kernel, representing the developmental progression. (c) UMAP visualization of limb bud cells, colored by cell-state density, highlighting regions of high and low cell density along the developmental trajectory. (d-f) UMAP visualizations as in (a-c), respectively, displayed using X_umap1 and X_umap2 dimensions for spatial context. (g) Scatter plot juxtaposing FT-Kernel predicted pseudotime against true pseudotime. Root Mean Square Error (RMSE) and the number of genes utilized for regression are indicated, quantifying the accuracy of pseudotime prediction. (h) Heatmap depicting pseudotime- related genes identified by FT-Kernel in human limb bud development. Gene expression levels are scaled and visualized to highlight temporal dynamics. (i) Cell- stage density distribution plot illustrating cell density distribution along the pseudotime trajectory of limb bud development. Density contours highlight areas of high and low cell state density across pseudotime. (j) Fate rank plot ranking genes by their contribution to cell fate determination in limb bud development, as determined by FT- Kernel. Higher rank indicates greater influence on cell fate. (k) Spatial plots exhibiting the expression of COL2A1 overlaid on spatial coordinates (spatial1 and spatial2) of the limb bud tissue. COL2A1, encoding collagen type II alpha 1 chain, is essential for cartilage formation, particularly in developing digits. (l) Spatial plots illustrating the expression of IRX1 overlaid on spatial coordinates (spatial1 and spatial2) of the limb bud tissue. IRX1, Iroquois homeobox 1, plays a critical role in limb patterning and digit identity. (m) Pseudotime dynamics of COL2A1 expression, depicting gene expression levels along the inferred pseudotime trajectory. The trend line represents the smoothed expression pattern. (n) Pseudotime dynamics of IRX1 expression, depicting gene expression levels along the inferred pseudotime trajectory. The trend line represents the smoothed expression pattern.

Furthermore, FT-Kernel robustly identified a discrete set of genes significantly associated with limb bud development (Fig. 6h). To investigate the spatial relevance of these identified genes, we spatially mapped the expression patterns of *COL2A1* and *IRX1*, two high-ranking fate determinants prioritized by FT-Kernel. Spatial coordinate plots revealed spatially complementary expression domains for *COL2A1* and *IRX1* (Fig. 6k, l). *COL2A1*, encoding collagen type II alpha 1 chain, a principal component of the cartilage extracellular matrix, exhibited pronounced expression within the ‘Digit’ region, concordant with its established and pivotal role in digit chondrogenesis and skeletal development. Conversely, *IRX1*, a member of the Iroquois homeobox gene family, known for its regulatory functions in limb anterior/posterior axis patterning and digit identity specification during limbogenesis, displayed elevated expression in the ‘HandPlate’ and ‘IDS’ regions bordering the ‘Digit’ domain. This spatial expression pattern of *IRX1* suggests its involvement in defining the spatial boundaries of the ‘Digit’ region and modulating cell fate in adjacent compartments. Temporal expression dynamics along the FT-Kernel inferred pseudotime trajectory further delineated distinct temporal regulatory modes for these fate determinants (Fig. 6m, n). *COL2A1* expression demonstrated a sustained increase along pseudotime, mirroring the progressive maturation of chondrocytes during digit development. *IRX1* expression, in contrast, peaked at early pseudotime and subsequently declined, implicating a critical role in early limb bud patterning and initial fate specification events. Furthermore, we observed a clear segregation of cell stage densities along the pseudotime trajectory (Fig. 6i), reinforcing the efficacy of FT-Kernel in capturing density-dependent fate transitions during limb bud development. Fate rank analysis further prioritized genes critically involved in cellular fate determination during limbogenesis (Fig. 6j). Collectively, these findings robustly demonstrate FT-Kernel as a potent analytical tool for dissecting the intricate spatiotemporal dynamics of cell fate decisions in complex developmental processes, readily applicable to spatially resolved transcriptomics datasets.

## Discussion

We have presented a new computational kernel, FT-Kernel, which effectively identifies cell-state density lineage transition factors along the pseudotime critical for cellular fate determination using scRNA-seq data. To our knowledge, FT-Kernel is the first tool to: (1) model cell-cell interaction dynamics along pseudotime (Fig. 4), and (2) integrate density-aware regression for gene selection (Fig. 2e-h). FT-Kernel overcomes significant limitations of traditional pseudotime-based gene identification methods by incorporating both temporal and density-based state transitions. Utilizing ridge regression and adaptive threshold regression (ATR), FT-Kernel isolates a minimal set of time-related genes that maintain pseudotime regression model adequacy, thus extracting the time-related fate-determining genes, cellular interactions, and genesets/pathways for the determined lineages from the cell-state density space. Furthermore, FT-Kernel is designed for scalability, making it applicable to large-scale datasets using GPU-accerlate.

The selection of performance metrics was guided by the inherent challenges of pseudotime analysis and the distinctive strengths of FT-Kernel. Traditional metrics like Pearson correlation, while prevalent, exhibit limitations in capturing non-linear relationships and suffer from “maxima/minima bias” (Fig. 2c), leading to the omission of crucial transitional state genes. Our chosen metrics are designed to overcome these limitations and comprehensively evaluate FT-Kernel. R²/MAE guarantee comparable regression accuracy to traditional methods. Kendall’s τ validates temporal ordering consistency. Importantly, Number of Genes and Linregress slope are novel metrics specifically designed to evaluate FT-Kernel’s key innovations: the efficiency of identifying minimal gene sets and the enhanced ability to capture genes crucial for intermediate state transitions. Number of Genes reflects the model’s efficiency in minimizing noise and experimental costs. Linregress slope directly addresses the limitations of Pearson correlation by specifically assessing the linearity of gene expression changes and enabling the identification of intermediate state genes often missed by traditional methods. Collectively, this metric panel provides a nuanced and robust evaluation of FT-Kernel, highlighting its advantages in deciphering the complex dynamics of cell fate determination along pseudotime.

Our study highlights the importance of cell-state density in understanding differentiation trajectories and demonstrates the potential of FT-Kernel to provide new insights into regulatory mechanisms that guide these processes along the pseudotime. We validated FT-Kernel’s efficacy across various datasets, including human hematopoiesis, mouse DG neuron development, and the decidua-placenta interface. The results demonstrate that FT-Kernel can accurately identify cell-state density lineage transition genes and reveal critical cellular interactions during lineage transitions. For instance, in hematopoietic differentiation, FT-Kernel identified key regulatory genes such as *EBF1* for B-cell development and *SOX6* for erythroid development, showcasing its capability to pinpoint time-related genes crucial for early lineage development.

Several important future extensions for FT-Kernel are envisioned. First, while our method currently focuses on scRNA-seq data, it can be extended to integrate other data modalities such as scATAC-seq or spatial transcriptomics, which provide additional layers of regulatory information. Second, incorporating machine learning techniques, including deep learning, could further enhance the identification of subtle gene expression patterns and improve the robustness of our predictions. Third, expanding FT-Kernel to include more complex cellular interactions and feedback loops could provide a more comprehensive view of cellular dynamics during development. FT-Kernel Linregress scales as (𝑂(𝑛𝑙𝑜𝑔𝑛) (n=cells). Benchmarks: 10k cells (1s), 100k cells (10s) on 16-core CPU. To thoroughly evaluate the practical applicability of FT- Kernel, we conducted a benchmark analysis on its scalability and computational efficiency, comparing it against other relevant algorithms (Fig. S2).

In conclusion, FT-Kernel offers a powerful and scalable tool for unraveling the complexities of cellular fate factors by integrating cell-state density and temporal dynamics. Our method not only improves the identification of key time-related regulatory genes but also provides a comprehensive framework for understanding the intricate processes governing the whole cellular developmental process. FT-Kernel is available as an open-source package, facilitating its adoption and further development by the research community.

## Materials and Methods

### Key resources table

**Table 1.**
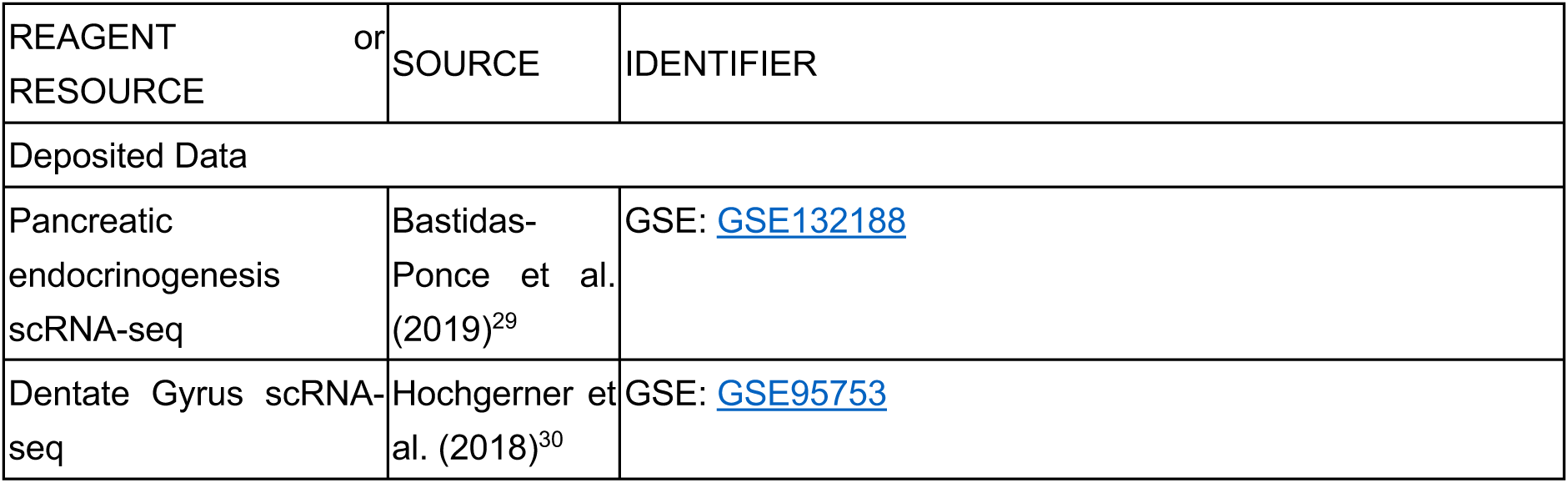
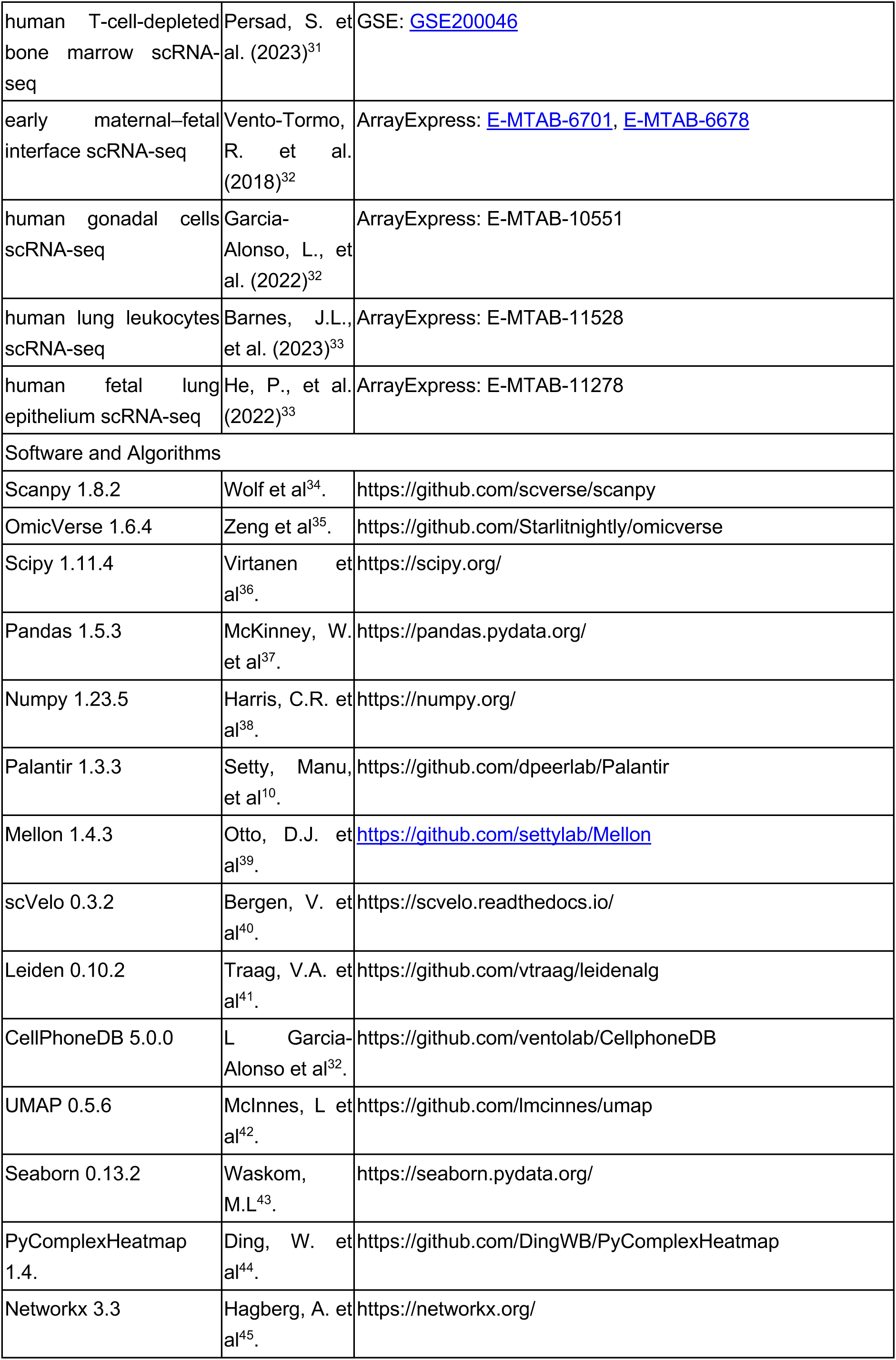
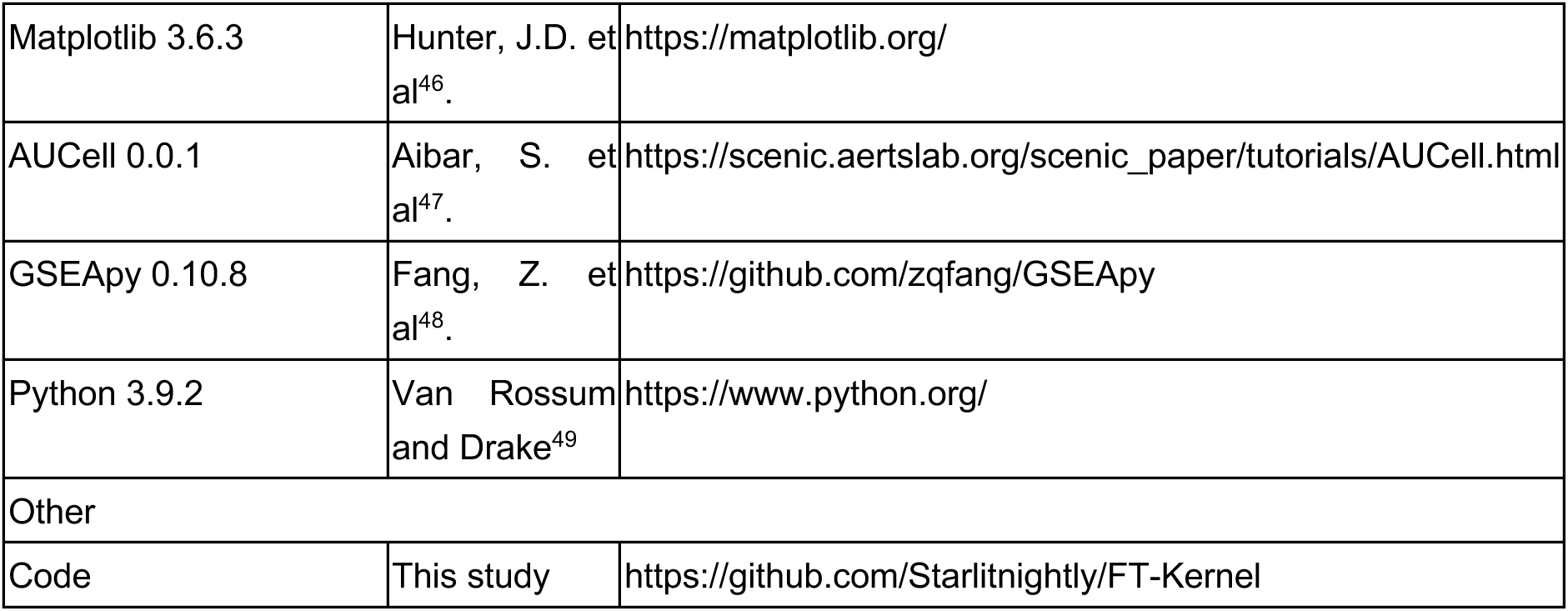
The version of the software used in this study.

### FT-Kernel Linregress algorithm

1. Gene/Peak Representation: Let 𝐺_!_denote each gene or peak. For various gene/peak combinations within each cell, we utilize gene sets (pathways, transcription factors (TF), and their targets) or cellular interactions. In the case of gene sets, we employ AUCell to compute the fraction of each gene set, which then substitutes for the individual gene/peak 𝐺_!_. For cellular interactions, CellPhoneDB is utilized to determine the receptor of the target cell. This receptor is then mapped to the gene set of the ligand cell, effectively replacing the gene/peak 𝐺_!_.
2. Model Definition: Let 𝐗 represent the matrix of different gene/peak combinations 𝐺_!_, and 𝐲 denote the vector of proposed chronological scores of the cells. We construct a gene set-proposed chronological model using Ridge linear regression, defined by:

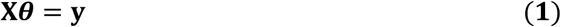

The loss function 𝐿(𝜽) for Ridge regression is given by:

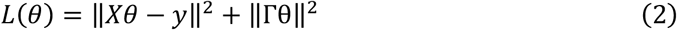

Here, ‖𝐗𝜽 − 𝐲‖” represents the squared errors between the predicted and actual values, and‖Γ𝜽‖ is the squared regularization term, and where Γ is selected via 5-fold cross-validation. We define Γ = 𝛼𝐈, where 𝐈 is the identity matrix, and 𝛼 is a regularization parameter.

1. Weight Dynamic Decay Model: To optimize the Ridge regression models for gene/peak combinations under different weight thresholds (denoted as 𝛼), we employ a weight dynamic decay model. This model evaluates changes in the mean squared error (MSE) of the regression coefficients and selects the target gene set combinations 𝐗_($%&)_ that maintain the minimum MSE for training parameters in the Ridge regression:

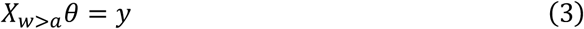

1. Parameter Calculation: Finally, we calculate the parameters 𝜽 of the Ridge regression model, which represent the weights of the genes/peaks within the regression model. Larger weights indicate gene sets with stronger associations to the proposed chronological scores, thereby providing valuable insights into the importance of different genes/peaks in the context of temporal differentiation.

This formalized approach ensures a rigorous and systematic analysis of the gene/peak associations with chronological scores, leveraging the strengths of Ridge regression and dynamic weight adjustment to uncover critical gene sets involved in cell differentiation trajectories. More detail of FT-Kernel Linregress algorithm algorithm could be found in supplement_pseudocode .

### FT-Kernel Branch Variability algorithm

For each cell 𝑖, the local gene expression variability 𝐿_!(_ is determined through the following steps: Neighborhood expression difference: We need to define the neighborhood 𝒩(𝑖) of cell 𝑖, Here, 𝒩(𝑖) = 0,1, …, 𝑁 − 1}, and 𝑁 denote the neighborhood of the cell 𝑖. Then we compute the expression difference Δ𝑋_!)(_ for each gene 𝑗 between cell 𝑖 and each cell 𝑘 ∈ 𝒩(𝑖):

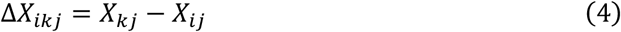

1. Rate of Euclidean Distance change calculation: We then Compute the Euclidean distance 𝑑_!)_ of the expression differences for each pair of cells (𝑖, 𝑘):

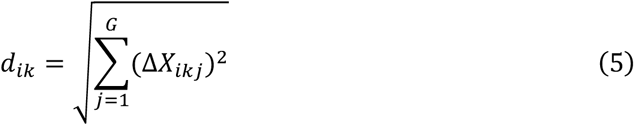

And the rate of euclidean Distance change 𝑅_!)(_ for each gene 𝑗 between cell 𝑖 and each cell 𝑘 ∈ 𝒩(𝑖):

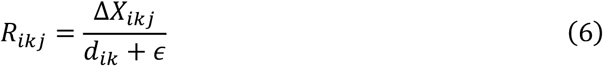

Here, ɛ be a small positive constant to prevent division by zero, typically set to 10^-,.^Local Variability Calculation: The local variability 𝐿_!(_ for each gene 𝑗 in cell 𝑖 is given by 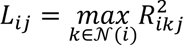, and the local variability matrix 𝐿 ∈ ℝ^1×*^ is constructed. Calculation of Gene Low-Density Variability Scores in Special branch: we use mellon to calculate the density 𝐷_!_ for each cell 𝑖, and we define the low-density weights 𝑊 based on the density 𝐷:

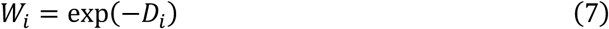

Then we mask 𝑀 to select cells of interest, 𝑀 denote the cell mask matrix of size 𝑁 × 𝐵, where 𝐵 represents the number of different branches or transitional states, and 𝑀_!3_ is 1 if cell 𝑖 belongs to branch 𝑏, other wise 0. We then selected the special branch 𝑏, and use the mask 𝑀 to select the indices ℐ_3_ of cells belonging the branch 𝑏. We multiply the local variability 𝐿_!(_ of these cells by their corresponding weights 𝑊_!_, and calculate the weighted average of the local variability for gene 𝑗 across all selected cells 𝑖:

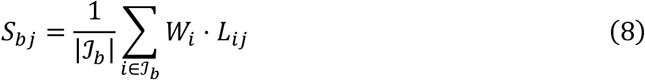

This formula indicates that the low-density variability score 𝑆_3(_ for gene 𝑗 in branch 𝑏 is the weighted average of the local variability for gene *j* across all cells in the branch, with weights emphasizing low-density cells. More detail of FT-Kernel Branch Variability algorithm could be found in supplement_pseudocode.

### scRNA-seq data preprocessing and analysis

For the preprocessing and analysis of single-cell RNA sequencing (scRNA-seq) data across various datasets, a standardized procedure was employed unless otherwise specified:

1. **Normalization**: Raw counts were normalized by dividing them by the total counts per cell to adjust for differences in sequencing depth.
2. **Log transformation**: The normalized data was multiplied by the median of total counts across cells and then log-transformed with a pseudocount of 0.1. This step aids in stabilizing variance and making the expression values more comparable between cells.
3. 3. **Feature selection**: The top 2500 most highly variable genes were selected from this transformed dataset, which served as input for principal component analysis (PCA) using 50 components.
4. Dimensionality reduction and clustering:

`o` PCs were utilized as inputs for performing leiden clustering to group cells into distinct cell types or states.
`o` UMAP (Uniform Manifold Approximation and Projection) visualizations were created from the PCA results to provide a low-dimensional embedding of the data that captures complex relationships between cells.
5. **Analysis**: The aforementioned procedures were carried out using the scanpy package, which offers comprehensive tools for single-cell analysis including normalization, clustering, and visualization.
6. Diffusion maps computation:

`o` Diffusion maps were computed using the Palantir2 package with default parameters on PCs as inputs.
`o` These maps provide a continuous representation of cell state transitions that can reveal underlying cellular differentiation pathways or states.
7. Gene expression imputation: The diffusion kernel was utilized for MAGIC8 gene expression imputation, which helps fill in missing values by leveraging the inferred cellular relationships.
8. Batch correction:

`o` Where applicable, batch correction using Harmony with default parameters was performed to adjust for technical variations between different sequencing batches.
`o` Batch-corrected PCs were then used as inputs for subsequent analyses such as UMAPs, diffusion maps, and gene expression imputation to ensure that the results are not biased by technical artifacts.

This rigorous methodology ensures that the analysis of scRNA-seq data is systematic, reliable, and capable of uncovering meaningful biological insights.

### Trajectory inference

To determine the pseudotime, or developmental timeline, of single-cell RNA sequencing (scRNA-seq) data from hematopoietic processes, we establish ’HSC’ as the initial stage (’origin’) and ’Naïve B’, ’Mono’, and ’Ery’ as terminal stages. This hierarchical model is then utilized by palantir for pseudotime calculation.

For pancreatic and dentate gyrus data analysis, we first compute the Mu/Ms matrix from the unspliced/spliced matrix using scVelo.pp.moment, which helps in identifying the momentary states of cells. Following this, dynamic models are estimated via scVelo.tl.dynamic_model. The pseudotime is then calculated for each cell with scVelo.tl.latent_time, allowing us to construct a trajectory that represents the developmental progression across these cellular datasets.

This comprehensive approach enables a detailed understanding and mapping of cellular differentiation processes across various biological systems.

### Pseudotime-related gene using pearson model

To quantify the relationship between gene expression levels and pseudotime along the trajectory of cellular development, we apply scipy.stats.pearsonr, a statistical method for assessing linear correlation between data points. A threshold of 70% of the maximum value is utilized to filter genes that are significantly correlated with pseudotime, thereby identifying those considered ’pseudotime-related’ using the Pearson model.

### Pseudotime-related gene using gene likelihood of dynamic model

To determine genes associated with pseudotime based on their likelihood under a dynamic model framework, we first fit all high variance genes as input data and then estimate dynamic models through scVelo. Subsequently, the gene likelihood scores for these high variance genes (HVGs) are computed.

### Protein Structure Alignment Methods

In this study, we employed protein structure alignment methods to analyze the structural relationship between ATP8B4 and ATP11C proteins. The structural model of ATP8B4 was derived from the AlphaFold Protein Structure Database (AlphaFold DB, version [Insert version number you used, e.g., Version 4]), UniProtKB accession number Q8TF62, accessible via https://alphafold.ebi.ac.uk/entry/Q8TF62. The experimental structure of ATP11C was obtained from the RCSB Protein Data Bank (Research Collaboratory for Structural Bioinformatics Protein Data Bank, RCSB PDB), PDB ID 7BSP, available for download at https://www.rcsb.org/structure/7BSP.

For structural comparison, we utilized the US-align web server (https://zhanggroup.org/US-align/). US-align is a widely used tool for protein structure alignment, providing reliable assessments of structural similarity. We focused on two key metrics from the US-align output: TM-score and RMSD. TM-score provides a quantitative measure of structural topology similarity, ranging from 0 to 1, with higher values indicating greater similarity. RMSD (Root Mean Square Deviation) reflects the average spatial deviation between the two aligned structures, with lower values indicating closer structural proximity. The resulting TM-score and RMSD values were used to quantify the structural similarity between ATP8B4 and ATP11C.

### Gene sets score calculated

To ascertain the score for each cell’s genesets, we start by obtaining pathway genesets from the enrichr database and preparing them using the omicverse.utils.pathway_prepare function. The AUCell scores for these genesets within individual cells are then calculated utilizing omicverse.utils.geneset_aucell. It is important to note that the pseudotime of each cell in this process is derived from its expression profile as captured by single-cell RNA sequencing (scRNA-seq).

### Cell-cell interaction analysis

To derive the ligand-receptor pairs for each individual cell from single-cell RNA sequencing (scRNA-seq) data, we initiate by isolating and standardizing the raw data to retain only essential information such as cell names, their types, and gene expressions, while discarding extraneous details. Subsequently, employing a statistical method tailored for cpdb_statistical_analysis, we compute the ligand-receptor pairs for each cell.

In this context, cellular receptor sets are conceptualized as a group of genes on which one cell’s ligands act upon another cell. This notion parallels that of pathways but is applied specifically to inter-cellular interactions. By leveraging these receptor sets, we can pinpoint external cells that contribute to the evolution or differentiation of specific cellular lineages.

### Trend of pseudotime-related Gene

We can visualize the pseudotime-related gene using heatmap, but we need to evaluate the effect of different model from the trend of pseudotime-related gene. Here, we propose a series of methods to evaluate. First, we sorted the cell along pseudotime 𝑡, and defined 𝑋 as expression matrix of 𝑛 gene in 𝑚 cells, 𝑋 ∈ ℝ^5×6^. And we can define the sorted 𝑋 as 𝑋_789:;<_ = 𝑋[:, 𝑎𝑟𝑔𝑠𝑜𝑟𝑡(𝑡)].

Next, we calculated the average index position for each gene 𝑖 (each row in matrix 𝑋_789:;<_), find the indices where its expression value exceed 80% of their maximum value and calculate the average index of these positions. We have

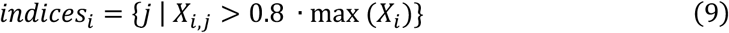

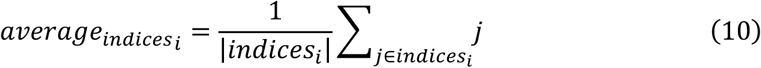

Store all average index values fro each gene into a list 𝐴:

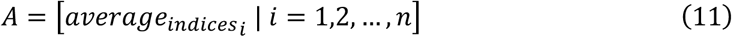

Finally, we calculate the Kendall Rank Correlation coefficient 𝜏 measures the relationship between two variables. With position indices 𝜏 = [1,2, …, 𝑛], calculate using ‘scipy.stats.kendalltau ‘:

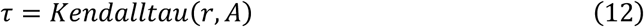

We also fit the relationship between 𝐴 and position indices 𝜏. The linear regression model can be represented as:

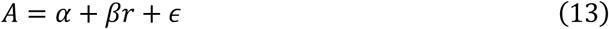

Where 𝛼 is the intercept, 𝛽 is the slope, and 𝜖 represents error terms. The result of linear regression can be found through using ‘scipy.stats.linregress’:

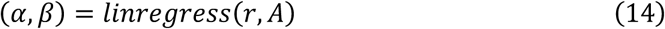

## Acknowledgments

Not applicable.

## Funding

This work was supported by the grants from the National Natural Science Foundation of China (32300682 to C.X.), the Fundamental Research Funds for the Central Universities (FRF-TP-22-007A1 to C.X.), the Student Research Training Program (SRTP) of University of Science and Technology Beijing (202010008107 to Z.Z.).

## Data and code availability

Data sets for hematopoietic stem cell differentiation are available on the Gene Expression Omnibus GSE200046^50^, for mouse pancreatic differentiation^51^ and dentate gyrus neuron differentiation^30^ are available from NCBI’s GSE132188 and GSE95753, and for trophoblast differentiation with cell classifications and the whole-genome sequencing data are deposited at ArrayExpress, with experiment codes E-MTAB-6701 (for droplet-based data), E-MTAB-6678 (for Smart-seq2 data) and E-MTAB-7304 (for the whole-genome sequencing data). All preprocessed data presented in this study is publicly available via the tutorial of FT-Kernel in omicverse. FT-Kernel (version: 1.0) is implemented as a Python package and is available through GitHub (https://github.com/Starlitnightly/TimeFactorKernel). Notebooks, tutorials for reproducing all figures in this study are also available through Github (https://github.com/Starlitnightly/TimeFactorKernel/tree/main/experiments)

## Conflict of interests

The authors declare that they have no competing interests.

## Author contributions

C.X., Y.M., and Z.Z. contributed equally. C.X. implemented the pipeline, performed the experiments, analysis, and wrote the manuscript. Z.Z. designed the algorithm of CellFateGenie, performed the benchmark test, and wrote the manuscript. Y.M. peformed the cell-cell interaction and pathway analysis, Y.W. and Y.W. polished the whole manuscript and tested the TimeFactorKernel. P.Q., H.D. and Z.Z. supervised and conceived the concept of TimeFactorKernel.

## Supplementary Figures

**Fig. S1.**
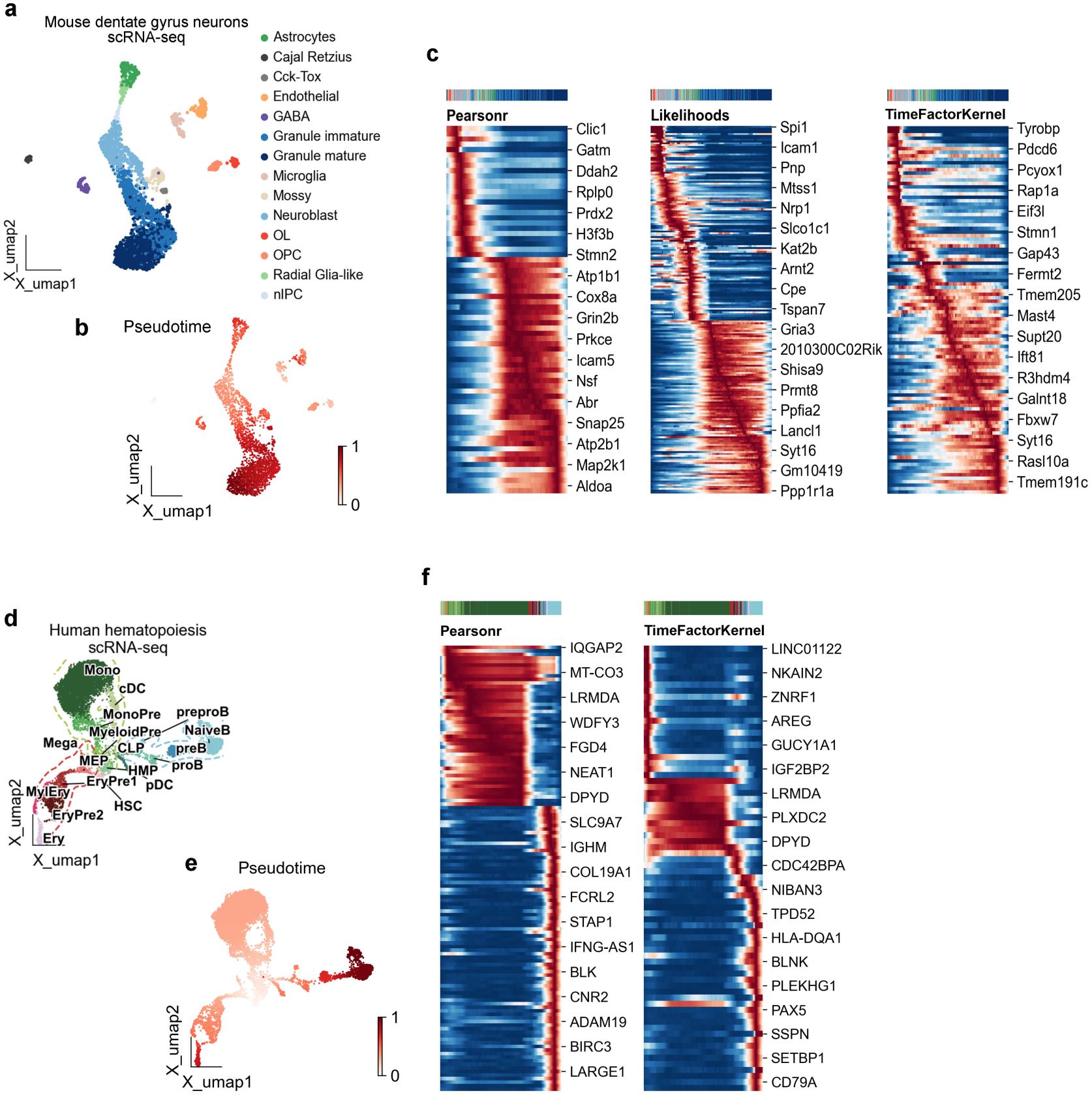
Kernel comparison using mouse dentate gyrus neurons and human hematopoietic stem cells scRNA-seq datasets. (a) UMAP of mouse dentate gyrus neurons scRNA-seq data, colored by cell types. (b) Same as a, colored by pseudotime calculated by scVelo. (c) Heatmap of pseudotime related genes, calculated by Pearsonr, Likelihoods, and TimeFactorKernel respectively. (d) UMAP of human hematopoietic stem cells scRNA-seq data, colored by cell types. (e) Same as d, colored by pseudotime calculated by Palantir for non-velocity-based data. (f) Heatmap of pseudotime related genes, calculated by Pearsonr, and TimeFactorKernel respectively.

**Fig. S2.**
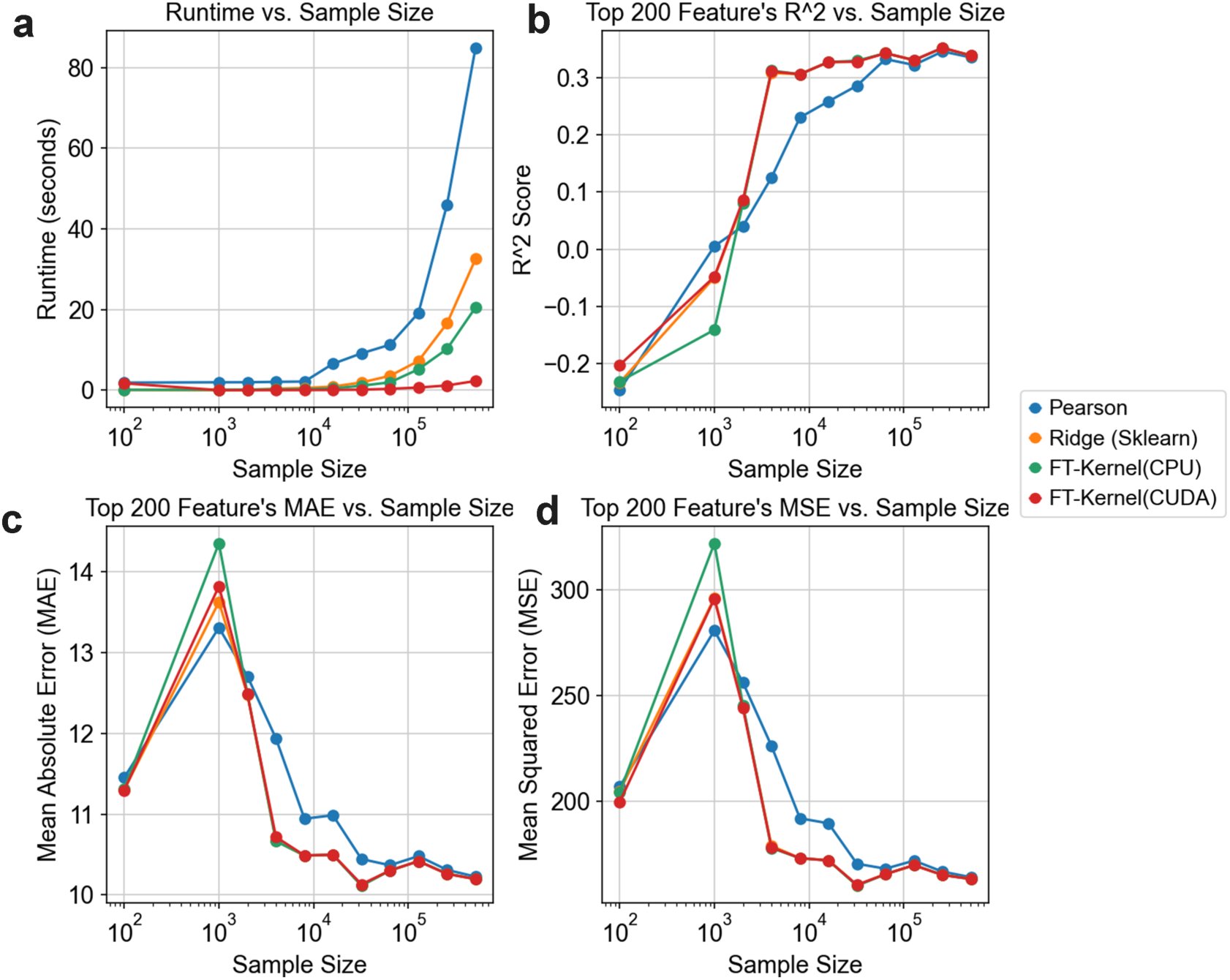
Evaluation of the scalability and computational efficiency of FT-Kernel. **(a)** Runtime comparison of different methods as a function of sample size (number of cells). FT-Kernel(CUDA) (red line) exhibits sublinear scaling and consistently lower runtime compared to other methods (blue: Pearsonr, green: FT- Kernel(CPU), orange lines (Ridge(Sklearn))), particularly with increasing sample size. The x-axis represents sample size on a logarithmic scale, and the y-axis shows runtime in seconds. **(b-d)** Performance metrics for different methods as a function of sample size, evaluated using the top 200 features identified by each method. **(b)** R-squared (R²) score, **(c)** Mean Absolute Error (MAE), and **(d)** Mean Squared Error (MSE). These plots demonstrate that while FT-Kernel maintains comparable performance in terms of R², MAE, and MSE to other methods, it achieves significantly better runtime scaling, as shown in panel **(a)**. The performance metrics are calculated based on a regression task using the top 200 features for each sample size. Different colors represent different methods, consistent across all panels.

## Notes

### Competing Interest Statement

The authors have declared no competing interest.

### Summary of Updates

Correct the name of author 2

